# Unraveling the stepwise maturation of the yeast telomerase

**DOI:** 10.1101/2021.04.30.442090

**Authors:** Anna Greta Hirsch, Daniel Becker, Jan-Philipp Lamping, Heike Krebber

## Abstract

Telomerases elongate the ends of chromosomes required for cell immortality through their reverse transcriptase activity. By using the model organism *Saccharomyces cerevisiae* we defined the order in which the holoenzyme matures. First, a longer precursor of the telomerase RNA, *TLC1* is transcribed and exported into the cytoplasm, where it associates with the protecting Sm-ring, the Est- and the Pop-proteins. This partly matured telomerase is re-import into the nucleus via Mtr10 and a novel *TLC1*-import factor, the karyopherin Cse1. Remarkably, while mutations in all known transport factors result in short telomere ends, mutation in *CSE1* bypasses this defect and become Type I like survivors. Interestingly, both import receptors contact the Sm-ring for nuclear import, which therefore resembles a quality control step in the maturation process of the telomerase. The re-imported immature *TLC1* is finally trimmed into the ~1150 nucleotide long mature form. TMG-capping of *TLC1* finalizes maturation, leading to mature telomerase.

## Introduction

The protection and maintenance of the ends of linear chromosomes during replication is crucial for dividing eukaryotic cells. Therefore, nucleoprotein structures called telomeres shield the ends of linear chromosomes from double strand break repair activities and thus from end-to-end ligations that would cause massive genome rearrangements and genome instability ^1, 2^. These telomeres are the target of a ribonucleoprotein (RNP) complex, called telomerase that protects the chromosome ends from progressive shortening by extending them through its reverse transcriptase activity. While in most somatic cells telomere shortening occurs after sufficient cell divisions and causes a cell to enter into replicative senescence, stem cells, germ cells, lymphocytes and unicellular eukaryotes like yeast express telomerase to encounter this effect. Remarkably, almost 90% of all cancer cells have re-activated the telomerase activity to bypass their proliferation limit ^1, 3, 4, 5^. As the production of functional telomerase is crucial for telomere maintenance, it is important to understand its assembly and the order of its step-wise maturation process to realize the underlying control mechanisms that generate functional telomerase.

The telomerase is an RNP complex that contains a long non-coding RNA component, *TLC1* (<u>te</u>lomerase <u>c</u>omponent 1) composed of 1158 nucleotides in yeast, which functions as template for the synthetic telomere repeats and as a scaffold for the protein components of the holoenzyme, providing the reverse transcriptase activity ^2, 6, 7, 8^ Like mRNAs and most lncRNAs, *TLC1* is transcribed in the nucleus by RNA polymerase II (RNAP II) and it is subsequently capped with a monomethyl guanosine (m^7^G) cap and polyadenylated like mRNAs ^6, 9^. However, in contrast to mRNAs, but very similar to the snRNAs, Tgs1 generates a 2,2,7-trimethylguanosine (TMG)-cap on *TLC1* in the nucleolus, which persists in the mature telomerase ^6, 10, 11^.

Interestingly, *TLC1* was described to exist in two forms in the cell, an 1158 nucleotide long mature form and a ~1300 nucleotide long precursor. The majority of the *TLC1* molecules that are present in cells is the short form and only ~10% of the *TLC1* RNA is the long polyadenylated form ^12^. This might reflect that approximately 10% of the *TLC1* RNAs might be in their maturation phase and 90% are already matured. However, in addition to the polyadenylation factor mediated transcription termination (CPF-CF), as used for mRNAs, a study uncovered the ability of the Nrd1-Nab3-Sen1 (NNS) system to terminate *TLC1* transcription and it was suggested that this would immediately generate the ~1150 nucleotides short form, characteristic of the mature telomerase ^13, 14^. If really both termination sites are equally used or if one is preferred over the other under certain conditions is currently unclear. Regardless of the transcription termination pathway used, the primary transcript is always longer than the mature form. Clearly, some of the *TLC1* molecules receive a ~80 nucleotide long poly(A) tail after transcription, which is removed during the maturation process and the exosome trims the pre-*TLC1* RNA to 1158 nucleotides ^6^.

In addition to these RNA-specific structural maturation events, *TLC1* provides a scaffold for several proteins ^15^. In analogy to the snRNAs, the 3’-end of *TLC1* is bound by a heptameric ring, composed of seven Sm-proteins, that encircles and stabilizes *TLC1* ^11, 16, 17^. Additionally, structurally important proteins as well as regulatory and catalytic factors bind to the *TLC1* RNA. One of the stem loops that are formed by *TLC1* is bound by the heterodimer Yku70-Yku80, which is important for a persisting nuclear localization through attachment of the telomerase to chromosome ends ^18, 19^. The central domain of *TLC1,* which includes the template domain for reverse transcription, is bound by the catalytic protein subunit Est2 and two accessory factors Est1 and Est3 ^2, 6^. The Est-proteins are stabilized by the Pop1, Pop6 and Pop7 proteins, which are also central components in the RNase MRP and the RNase P ^20, 21, 22^.

Besides *TLC1* processing and the protein loading onto this scaffold, this RNA undergoes nucleo-cytoplasmic shuttling ^6, 23^. It is exported into the cytoplasm via Mex67-Mtr2 and Xpo1/Crm1 ^23^. On mRNAs and snRNAs the RNA-contact of Mex67 is mediated by the guard proteins Npl3, Gbp2 and Hrb1 ^24, 25, 26, 27^. Thus, a similar mechanism is conceivable for *TLC1*. For Xpo1/Crm1 a physical RNA contact was shown via the cap binding complex (CBC) that interacts with the monomethyl cap ^24^.

After loading of the Est-proteins and the Sm-ring in the cytoplasm, *TLC1* is re-imported into the nucleus via Mtr10 ^23, 28, 29^. Prevention of shuttling, either by mutations in the export factors or the import factor results in telomere shortening defects and reduced *TLC1* levels ^23, 29, 30^.

For snRNAs the transfer to the cytoplasm was shown to be crucial for the generation of a functional spliceosome, because when not exported, the longer, unprocessed pre-snRNAs can be incorporated into the spliceosome and jeopardize splicing ^24^. Thus, the immediate cytoplasmic drain of these noncoding RNAs is crucial for cell survival and might also be similarly important for the transcribed immature *TLC1*.

Interestingly, maturation events such as trimming and TMG-capping were suggested to be nuclear events that occur prior to *TLC1* shuttling and current models suggest that only the Est-proteins and the Sm-ring are loaded in the cytoplasm ^23, 29,30^. Therefore, we investigated the maturation steps of *TLC1* in detail and uncovered a stepwise process. We found that pre-*TLC1* shuttles into the cytoplasm for the loading of the Sm-ring, the Est- and the Pop-proteins. The subsequent re-import of the protein bound *TLC1* precursor not only requires Mtr10, but also Cse1, a novel import receptor for *TLC1.* Defects in *CSE1* result in Type I like survivors that show an increased recombination phenotype, which is based on telomere elongation with increased Y’ elements. Both importer contact *TLC1* via the Sm-ring. In this way the nuclear entry of immature, or incomplete assembled *TLC1* RNPs is prevented. Thus, nuclear re-import resembles a quality control step in the life cycle of the telomerase RNA. We further show that trimming of *TLC1* to the 1158 nucleotide short form is carried out by the nuclear exosome after shuttling. In a final step, TMG-capping of *TLC1* completes maturation. In summary we have uncovered that the maturation of *TLC1* occurs in a different order than anticipated and in a highly ordered manner, including a cytoplasmic quality control step to ensure proper telomere function.

## Results

### The Sm-ring is loaded to the precursor of *TLC1* in the cytoplasm

Sm-ring binding to *TLC1* is a prerequisite for proper processing of this non-coding RNA, while the contact of the TRAMP-complex and the nuclear exosome prior to the Sm-ring attachment rather initiates full degradation of the transcript ^16, 17^. On snRNAs, Sm-ring loading was recently identified to be a cytoplasmic event ^24^. It was shown that newly transcribed snRNAs are immediately exported from the nucleus to prevent an incorporation of the precursor RNAs into the spliceosome, which jeopardizes splicing ^24^. *TLC1* receives its Sm-ring also in the cytoplasm (Fig. 1 and ^29^). To analyze to which form the Sm-ring is loaded, the short 1158 nucleotide long mature form or the longer premature form, we did the following experiments. First, we extracted GFP-tagged Smb1 from cell lysates (Fig. 1A) and analyzed in RNA-co immunoprecipitation (RIP) experiments the general binding of *TLC1*. Subsequent qPCR experiments showed a clear binding of *TLC1* to Smb1, which was normalized to no tag as a negative control and related to 21S rRNA (Fig. 1B). Importantly, when we repeated the experiment in the export mutant *mex67-5 xpo1-1* after shifting the cells for 2h to the non-permissive temperature, which prevents the export of newly synthesized *TLC1* molecules, we detected a significantly decreased binding between the Sm-ring component Smb1 and *TLC1* (Fig. 1C, D). These results suggest that the inhibition of the nuclear export of *TLC1* prevented the loading of the Sm-ring in the cytoplasm.

**Figure 1.**
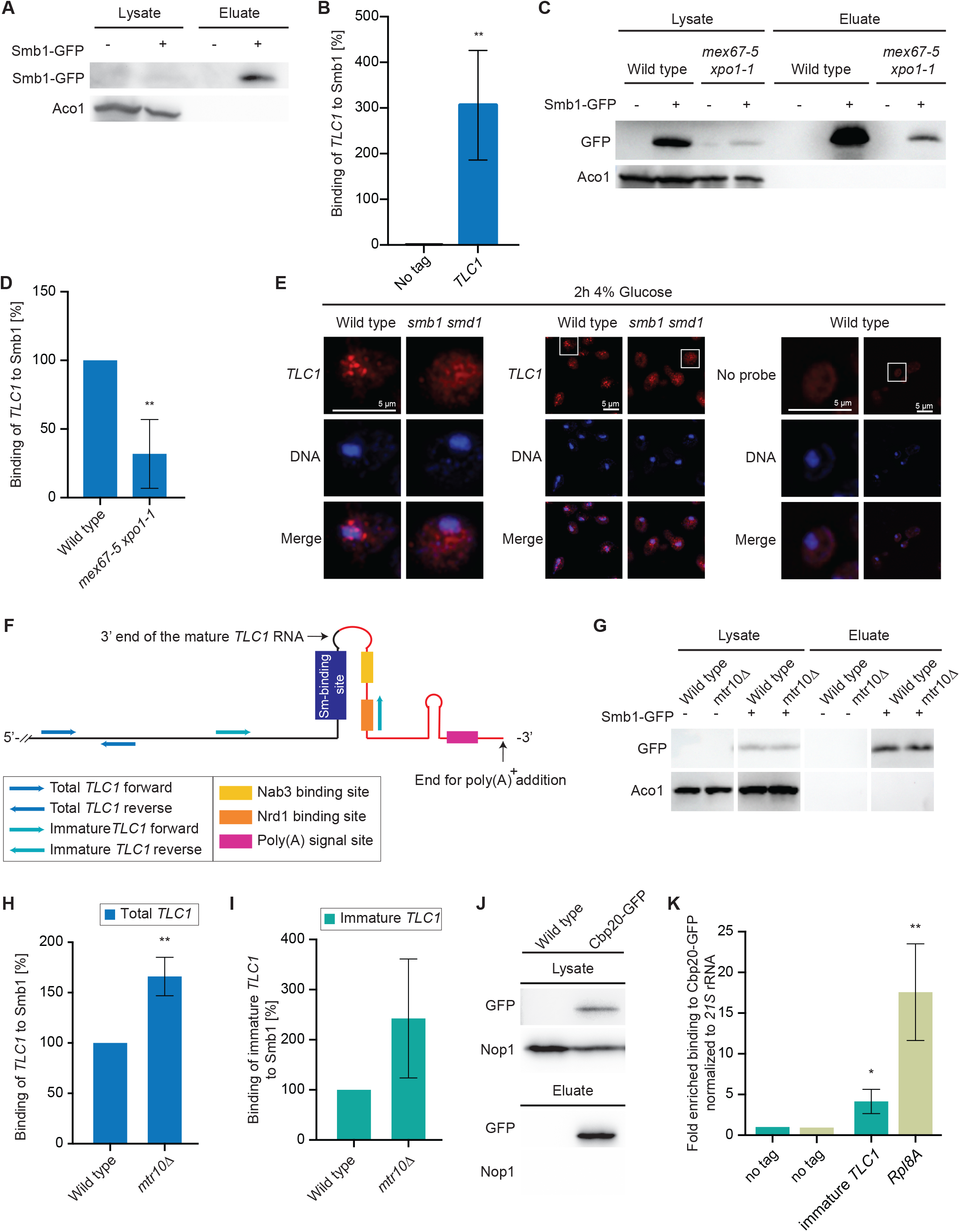
The longer precursor of *TLC1* binds the Sm-ring in the cytoplasm. (A) RIP experiments were carried out with GFP-tagged Smb1 from wild type cells. One example IP of Smb1 is shown on western blots. Aco1 served as a negative control. (B) Smb1 binds to *TLC1*. The Smb1 co-precipitated RNA was used as a template in qPCRs for *TLC1*. For this (and all following IPs carried out in this study), the amount of the eluted RNA was related to the RNA present in each lysate normalized to the mitochondrial 21S rRNA and compared to no tag. *n = 4*. (C) A western blot shows an example IP of the precipitated Smb1 protein from the RIP experiment shown in (D). (D) Smb1 cannot bind to *TLC1* when its nuclear export is prevented. The Smb1 co-precipitated RNA was used as a template in qPCRs for *TLC1* in the indicated strains, shifted to 37°C for 1h. *n = 3*. (E) *TLC1* accumulates in the cytoplasm of cells in which the Sm-ring is not properly assembled. FISH-experiment of a Cy3-labeled probe (red) targeting *TLC1* is shown in wild type cells and a strain in which the Sm-ring is not assembled trough mutation of *SMB1* and *SMD1* after glucose induced expression repression. The DNA is stained with DAPI. *n=3* (F) Scheme of the ~1300 nucleotide long immature *TLC1* and the primer positions for the amplification of the total and the immature *TLC1* molecules. (G) Western blot showing an example of a Smb1 IP used in Fig. 1H and I in RIP experiments. (H) Blocking nuclear re-import of *TLC1* increases the amount of the total *TLC1*. RIP experiments and subsequent qPCRs were carried out to show the total *TLC1* in the indicated strains. *n=3* (I) Blocking nuclear re-import of *TLC1* increases the amount of the immature *TLC1*. RIP-experiments were carried out and subsequent qPCR results of the immature *TLC1* in the indicated strains is shown. *n=3* (J) Western blot showing an example of a Cbp20 IP used in Fig. 1K for RIP experiments. (K) Immature *TLC1* binds to Cbp20. The Cbp20 co-precipitated RNA was used as a template in qPCRs for immature *TLC1. RPL8A* served as positive control. *n = 3*

This finding is further supported by a fluorescense *in situ* hybridization (FISH) experiment targeting the *TLC1* RNA with a Cy3-labeled probe. *TLC1* is mostly detectable in the nucleus of wild type cells (Fig. 1E) and ^23^. However, in a double mutant of *smb1 smd1*, which has a defective Sm-ring ^31^, *TLC1* mislocalizes to the cytoplasm (Fig. 1E). These findings confirm that Sm-ring loading to *TLC1* occurs in the cytoplasm, which was also shown earlier via an inducible *TLC1-*RNA tagging experiment and is very similar to the cytoplasmic Sm-ring loading onto the snRNAs ^24, 29^. Therefore, we investigated the interaction between the Sm-ring and *TLC1* in the import receptor mutant *mtr10*Δ. RIP-experiments showed an increased binding of total *TLC1* (Fig. 1G-I). To get information whether the mature or the immature form of *TLC1* accumulates, we also investigated the binding of the precursor of *TLC1* to Smb1. A scheme of *TLC1* shows that the primers amplifying the unprocessed variant of *TLC1* will amplify any generated pre-*TLC1*, either the through NNS or CPF-CF termination generated form (Fig. 1F).

Indeed, as shown in Fig. 1I, the binding of the immature *TLC1* to Smb1 increased even more than the total *TLC1*, suggesting that the immature form accumulates in *mtr10*Δ. These findings indicating that the Sm-ring is loaded in the cytoplasm to the immature pre-*TLC1* and the Sm-ring loaded *TLC1* is accumulating in the cytoplasm.

Futhermore, it was shown that the export of *TLC1* requires Xpo1 in addition to Mex67 ^23^. For snRNA export it was shown that Xpo1 contacts the RNA via the cap binding complex (CBC) ^24^. Since pre-*TLC1* contains an m^7^G cap, it seems likely that this 5’ cap is also bound by CBC. However, this has not been shown so far. Therefore, we carried out RIP experiments with Cbp20 and showed that the immature form of *TLC1* associates with this CBC-component (Fig. 1J-K). Together, these findings suggest that pre-*TLC1* export is supported by Xpo1.

### Cse1 is a novel nuclear re-import factor for *TLC1*

Nuclear re-import of *TLC1* requires the nuclear import receptor Mtr10 ^28^. Interestingly, snRNAs are re-imported via Mtr10 and Cse1, another member of the kayopherin transport factor family ^24^. After loading of the Sm-ring, Cse1 contacts the ring and supports nuclear re-import of the snRNA, which resembles an important control step for maturation, because the importer can only bind when the Sm-ring was successfully loaded ^24^. As the Sm-ring is also loaded onto *TLC1* in the cytoplasm, it seems reasonable to assume that Cse1 would also participate in its nuclear re-import. Therefore, we localized *TLC1* in the *cse1-1* mutant that was shifted to 16°C, as it is cold sensitive. We found that while *TLC1* localized mainly to the nucleus in wild type cells, it was distributed throughout the cytoplasm in the *cse1-1* mutant (Fig. 2A), similar to *mtr10*Δ and *yku70*Δ strains (Fig. 2A) and ^23, 28^. Since Mtr10 and Cse1 both contribute to the nuclear import of *TLC1* and snRNAs, we created a double mutant and assayed its growth. Although the single mutants already showed growth defects in comparison to wild type, the double mutant grew even less, indicating a genetic interaction between these import receptor genes (Fig. 2B). Subsequent FISH experiments with the double mutant also showed a cytoplasmic accumulation, similar to that observed in the single mutants (Fig. 2A). However, the overall signal in the double mutant was slightly less intense, which might reflect a decreased stability of *TLC1* in this strongly growth compromised strain.

**Figure 2.**
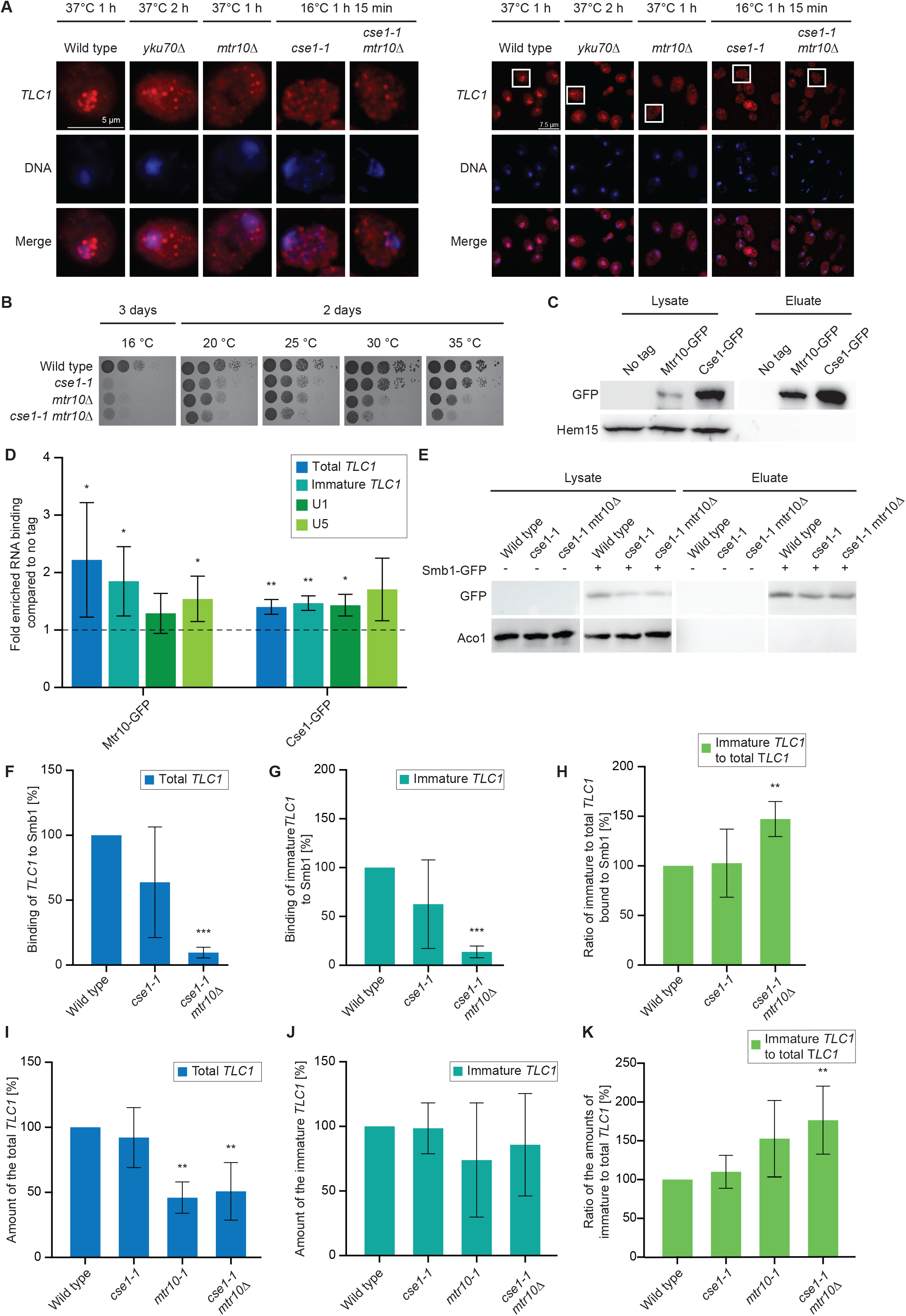
Cse1 is a novel nuclear import factor for *TLC1*. (A) *TLC1* mislocalizes to the cytoplasm in *cse1* mutants. A FISH-experiment for *TLC1* (red) is shown in the indicated strains that were shifted to the indicated non-permissive temperatures. *n= 3* (B) *CSE1* and *MTR10* genetically interact. Serial dilutions of the indicated strains were spotted onto full medium agar plates and incubated at the indicated temperatures. (C) RIP experiments were carried out with GFP-tagged Mtr10 and Cse1 from wild type cells. One example IP is shown on a western blot. Hem15 served as a negative control. (D) Cse1 binds to *TLC1*. The Cse1 co-precipitated RNA was used as a template in qPCRs for total *TLC1*. *n = 3*. Mtr10 served as a positive control. *n = 4*. (E) RIP-experiments were carried out in the indicated mutant strains. A western blot of an example IP is shown. Aco1 served as a negative control. (F) Binding of total *TLC1* to Smb1 decreases in *cse1-1* and *cse1-1 mtr10*Δ cells. RIP-experiments and subsequent qPCRs were carried out. *n=4* (G) Binding of immature *TLC1* to Smb1 increases in *cse1-1* and *cse1-1 mtr10*Δ cells. RIP-experiments and subsequent qPCRs were carried out. *n=4* (H) The ratio of the Smb1-bound immature versus total *TLC1* increases in *cse1-1 mtr10Δ.* The ratios were calculated from the experiments shown in F and G. *n=4* (I-K) The immature form of *TLC1* accumulates in nuclear re-import mutants. The RNA was isolated from the indicated strains after a 1h 15 min temperature shift to the non-permissive temperature. Subsequent qPCRs were carried out with primers that detect either all *TLC1* forms (total *TLC1*) (I) or only the unprocessed form (immature *TLC1*) (J). (K) The ratio of immature to total *TLC1* increases in the import mutants. The ratios were calculated from the experiments shown in A and B. *mtr10-1 n =3. cse1-1 n =6* and *cse1-1 mtr10Δ n =5*

To verify binding of *TLC1* to Cse1, we carried out RIP experiments, in which we purified GFP-tagged versions of the import receptors Cse1 and Mtr10, of which the latter served as a positive control (Fig. 2C, D). We found that the binding of *TLC1*, most likely its immature form was significantly enriched, similar to the binding of the snRNAs U1 and U5 (Fig. 2D) and ^24^. A 1,5 to 2-fold increase of the binding, although significant, might on first sight not seem to be very strong, but considering that the binding of both import receptors depends on the small GTPase Ran the low value is understandable. For import-cargo complex formation Ran must be in its GDP bound state. This is favored in the cytoplasm by the presence of the GTPase activating protein RanGAP1 and its co-factor RanBP1 ^32, 33, 34^. Cargo release from the import receptors occurs in the nucleus where RanGTP is present and the nucleotide exchange occurs with the nuclear exchange factor Prp20 (human RCC1) ^34^. Upon cell lysis for the RIP experiment, both compartments get mixed up and pre-formed nuclear import complexes are attacked and disassembled through RanGTP. An additional difficulty is the low copy number of *TLC1* with ~10 to ~30 copies per haploid cell ^6, 35, 36^. However, despite these difficulties, we were able to detect binding of *TLC1* to both import factors, Mtr10 and Cse1 (Fig. 2C, D).

Analogous to *mtr10*Δ (Fig. 1H, I) we tested the binding of the Sm-ring in *cse1-1* and the *cse1-1 mtr10*Δ double mutant. In contrast to *mtr10*Δ we see a rather decreased binding of total *TLC1* and unprocessed *TLC1* in *cse1-1*, although the ratio of immature *TLC1* still exceeds the total *TLC1* (Fig. 2 E-H). This observation suggests that Cse1 prevents degradation of *TLC1* when associated with the Sm-ring. In the double mutant, in which both import receptors are missing, the Sm-ring binding is even further reduced and the stability of *TLC1* is even worse, suggesting that both import factors contribute to stabilization of the RNA waiting to be imported. However, Cse1 seems to be important for the initial Sm-ring loading. In *mtr10*Δ the Sm-ring bound RNA is more stable, suggesting that Cse1 is already sufficient to protect the immature *TLC1* in the RNP. Remarkably, we still detect higher amounts of the immature *TLC1* bound to the Sm-ring (Fig. 2H), suggesting that this form is trapped in the cytoplasm.

Similar results were also obtained when we analyzed the complete *TLC1* content in cells. While the total *TLC1* level decreased to approximately 50% *in mtr10-1* and the double mutant *cse1-1 mtr10*Δ, the immature form was present in higher amounts (Fig. 2I-K), suggesting a constant production of pre-*TLC1*, which does not finish maturation. Together, these findings identify Cse1 as a novel import receptor for *TLC1* and show that optimal nuclear re-import of this a large RNP relies on more than one import receptor for its re-import and its stabilization.

### Mutation of *CSE1* generates a Type I like survivor phenotype

Prevention of nuclear *TLC1* shuttling leads to shortened telomeres and thus mutations in the export receptor Mex67 and Xpo1 as well as in the import receptor Mtr10 have been reported to result in telomere shortening defects ^23, 30^. To investigate whether such defects would also be visible in the new import factor Cse1, we carried out southern blot analyses. Strikingly, we detected no shorter telomere ends in *cse1-1*, but instead highly amplified subtelomeric Y’ elements (Fig. 5A, B). Usually cells that lack the telomerase or components of it undergo senescence when telomeres become critically short. The senescence phenotype is only temporary and most of these cells lose their viability, but a small percentage of cells overcome this phenotype by lengthening telomeres through homologous recombination, termed Type I and Type II survivors ^37, 38^. Type I survivors depend on Rad51 and exhibit highly amplified subtelomeric Y’ elements ^39^, which are clearly visible in *cse1-1* cells (Fig, 5B and 5C). Thus, while the export receptor mutants and *mtr10Δ* were not able to switch their telotype to become Type I or Type II survivors and maintain stable but short telomeres, mutation in *CSE1* allowed homologous recombination, resulting in amplified Y’ elements to create Type I survivors, even without a prior senescence phenotype. However, untypical for Type I survivors is that they have no short telomeres, as the case in *cse1-1* cells. Therefore, we designated them as Type I *like* survivors.

**Figure 5.**
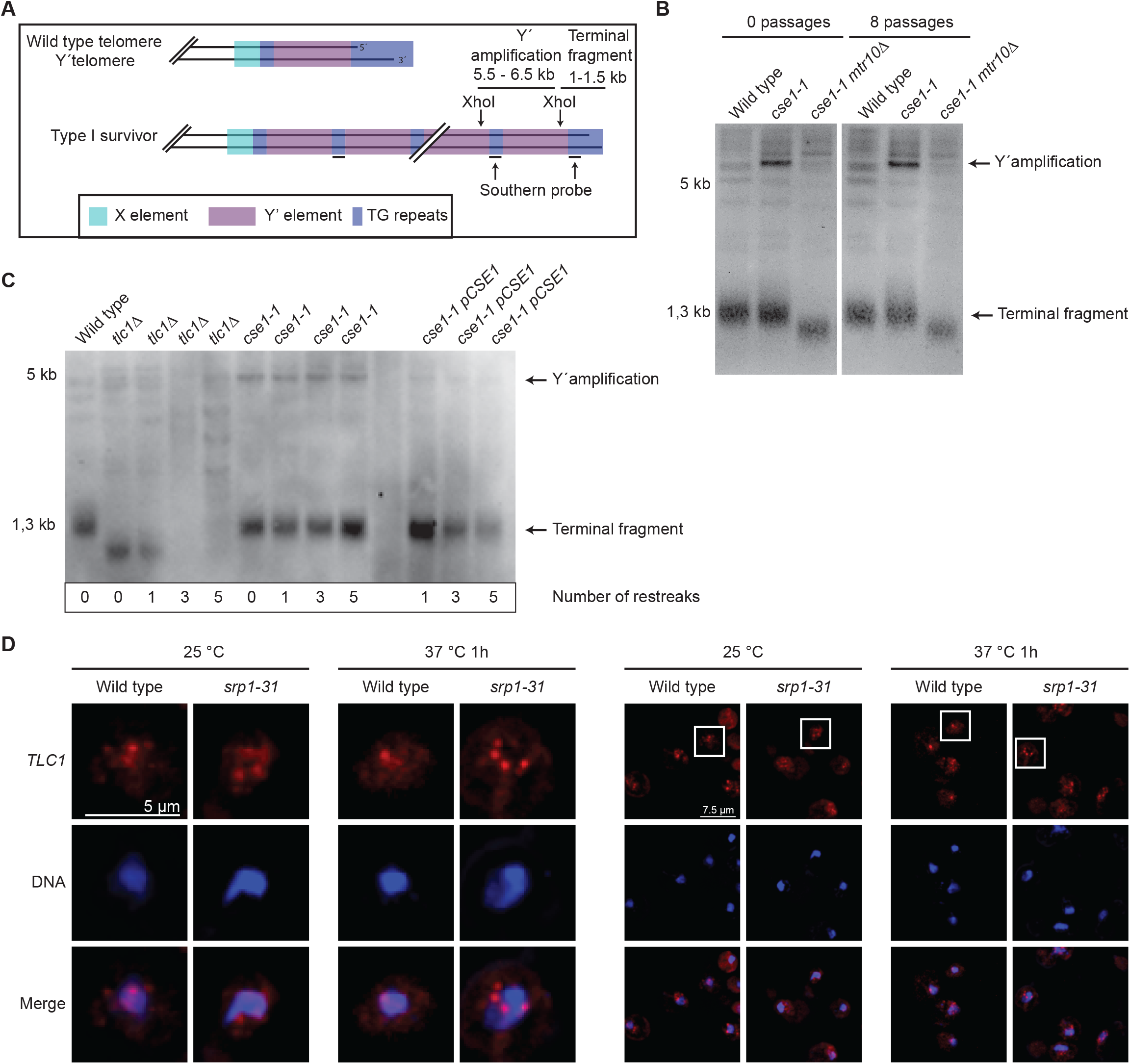
Mutation of *CSE1* results in a Type I like survivor phenotype. (A) Scheme of the Y-amplification and location of the probe that was used for the southern blot. (B) The *cse1-1* mutation leads to the amplification of the telomeric Y-element. XhoI digested genomic DNA of the indicated strains was used for the southern blot. The chromosome ends were detected with a digoxygenin labeled probe. (C) Plasmid encoded *CSE1* recues the *cse1-1* phenotype. XhoI digested genomic DNA of the indicated strains was used for the southern blot. The chromosome ends were detected with a digoxygenin labeled probe (Fig. 5A). (D) *TLC1* localization is not altered in the *srp1-31* mutant. FISH expermiments show no mislocalization of *TLC1* in *srp1-31* after shift to 37 °C for 1h. n=3.

In addition, we analyzed whether the telomeric phenotype in the *cse1-1* mutant can be reversed by introduction of a plasmid containing wildtype *CSE1*. Already after one restreak, which equals approximately 25 generations, and more clearly after 3 restreaks, the Y’ amplification band disapeared, indicating that the phenotype is based on the *cse1* mutation (Fig. 5C). Furthermore, it can be seen that upon restreaking of the *tlc1Δ* strain on a plate, Type I survivors are generated ^40^ and the typical Y’ band that appears is at the hight of that seen in *cse1-1* (Fig. 5C).

Cse1 is also involved in the recycling of importin alpha, encoded by *SRP1* and *cse1-1* leads to an accumulation of Srp1 in the nucleus ^41^. It was reported earlier that Srp1 might be involved in the nuclear localization of Est1 ^42^. Therefore, we examined whether *TLC1* mislocalizes in an *srp1-31* mutant. FISH experiments show no mislocalization of *TLC1* in *srp1-31* (Fig. 5D).

### Pop protein loading onto *TLC1* occurs in the cytoplasm

After knowing the nuclear re-import requirements for *TLC1*, we asked, where the proteins of the RNP are loaded onto *TLC1* in the cell. We have suspected earlier that Est-protein binding occurs in the cytoplasm, as the Est-proteins accumulate in this compartment in a *tlc1*Δ mutant ^23^. Reassuringly, we also detect a cytoplasmic mislocalization for Est1-GFP in the new *TLC1* re-import factor mutant *cse1-1* (Fig. 3A, B).

**Figure 3.**
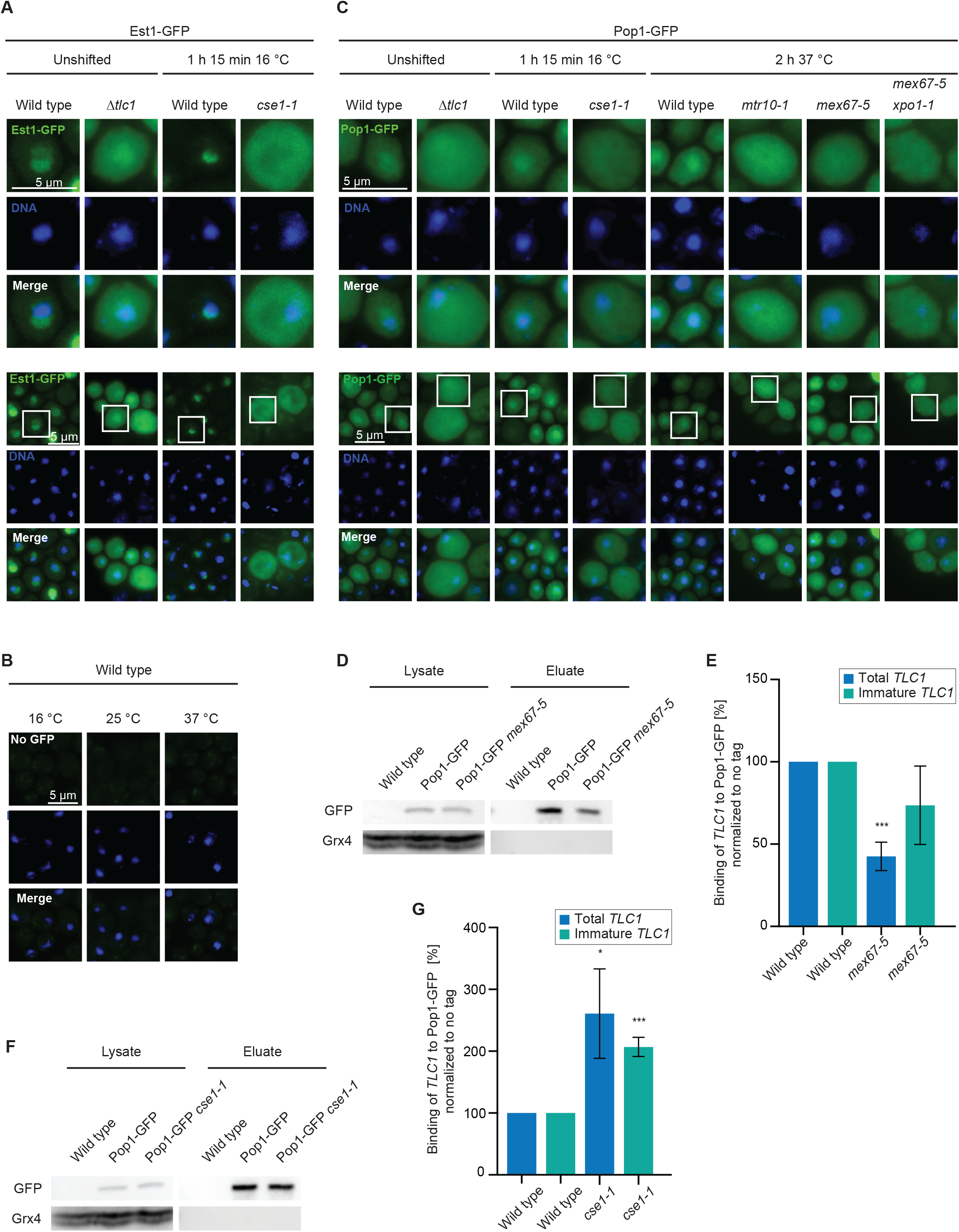
Loading of the Est and Pop proteins occurs in the cytoplasm. (A) Nuclear GFP-tagged Est1 mislocalizes to the cytoplasm in *cse1-1* cells. (B) A wild type strain is shown without a GFP tagged protein with identical microscopy settings as shown in (A). (C) Nuclear Pop1 localization is disturbed in mutants affecting *TLC1* expression or its localization. *n=3*. (D) Western blot of the Pop1-GFP IP is shown in the indicated strains. Grx4 served as a negative control. (E) Pop1 binding to *TLC1* is decreased in an export mutant. The Pop1-bound RNA from the several IP, one of which is shown in D, was analyzed in qPCRs *n = 3*. (F) Western blot of the Pop1-GFP IP is shown in the indicated strains. Grx4 served as a negative control. (G) Pop1 binding to *TLC1* is increased in an import mutant. The Pop1-bound RNA from the several IP, one of which is shown in F, was analyzed in qPCRs. *n = 3*.

For the Pop proteins the place of loading onto *TLC1* is currently unknown. As the proteins are mostly localized to the nucleolus and the nucleoplasm ^43^, it is possible that they are loaded after *TLC1* has completed its cytoplasmic phase and returned to the nucleus. The function of the Pop-proteins in *TLC1* maturation is to stabilize Est1 and Est2 within the holoenzyme ^21, 22^. Therefore, it is also conceivable that they are loaded to *TLC1* in the cytoplasm, after the Est proteins were properly attached. Importantly, Pop1 can only be stably associated with *TLC1* when the heterodimer Pop6 and Pop7 was attached to the RNA ^44^. Thus, Pop1 binding is the final step in the loading of the Pop-protein complex. We localized Pop1 in the *TLC1* nuclear import- and export mutants and in *tlc1*Δ and found a clear cytoplasmic mislocalization of the protein, even though we detect its localization in wild type not as strict nuclear as published earlier with the identical constructs ^43^ (Fig. 3C). Interestingly, also mutations in the Pop-proteins were shown to increase the cytoplasmic presence of *TLC1* ^20^, which supports a model in which loading of the Pop-proteins might occur in the cytoplasm. Thus, in situations, in which *TLC1* is not present in the cytoplasm, such as in *tlc1*Δ or in the export mutant *mex67-5 xpo1-1*, the Pop-proteins accumulate in the cytoplasm and most likely do not bind *TLC1*. However, in the import mutants, one would expect that the proteins were loaded onto *TLC1*, but cannot be imported. To get additional support for a cytoplasmic loading and to find out whether Pop1 is loaded onto the long immature *TLC1* we carried out RIP experiments, with strains that contained endogenously tagged *POP1-GFP* expressed from its own promoter to avoid overexpression. Clearly, we found a decreased binding of this RNA in the export mutant (Fig. 3D, E), while an import block resulted in an increased binding of the immature *TLC1* to Pop1 (Fig. 3F, G), suggesting that the loading of the Pop proteins is indeed a cytoplasmic event.

### Mtr10 contacts *TLC1* at the Sm-ring for nuclear re-import

Cse1 can mediate the nuclear import of snRNAs and *TLC1* only after the Sm-ring has properly assembled ^24^ and (Fig. 1E). An interaction between the Sm-ring and Cse1 has been shown before ^24^. Their interaction thus resembles a quality control step to allow only nuclear entrance of the Sm-ring-bound *TLC1* RNP. It seems possible that also Mtr10 may contact *TLC1* only after proper protein loading has occurred. To analyze potential complex formations, we carried out co-immunoprecipitations (co-IPs) of Mtr10 with Pop1 and Est1. However, none of these proteins physically interacted with Mtr10 (Fig. 4A and B).

**Figure 4.**
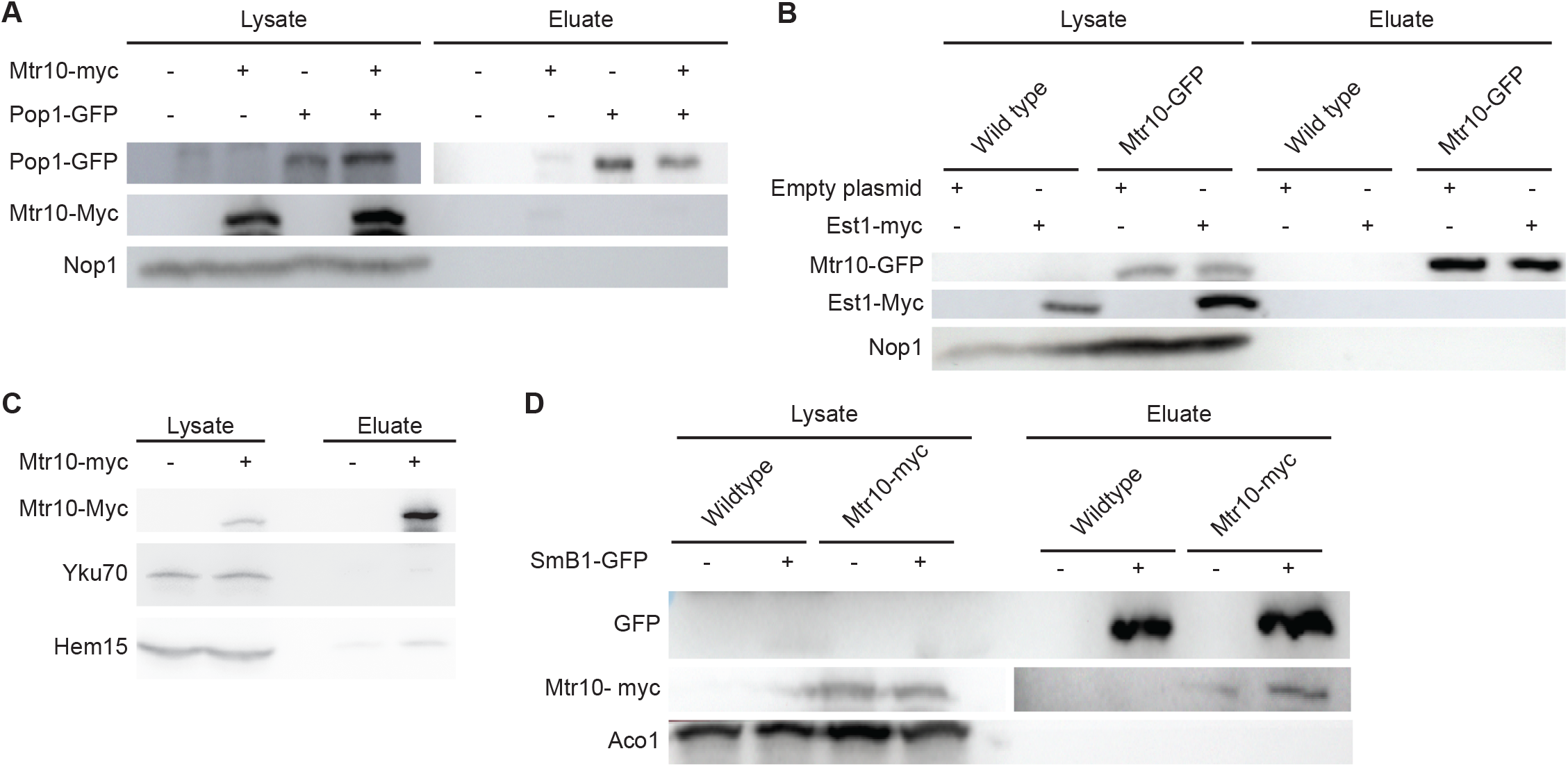
Both nuclear import receptors contact *TLC1* at the Sm-ring. (A) Mtr10 does not interact with the Pop-protein complex. A western blot of co-IPs of GFP-tagged Pop1 with myc-tagged Mtr10 is shown. Nop1 served as a washing control *n=3* (B) Mtr10 does not interact with the Est protein complex. A western blot of a co-IP of GFP-tagged Est1 with myc-tagged Mtr10 is shown. Hem15 served as negative control. *n=3* (C) Mtr10 does not interact with the Yku protein complex. A western blot of a co-IP of myc-tagged Mtr10 with Yku70 is shown. *n=3* (D) Mtr10 interacts with the Sm-ring. A western blot of a co-IP of GFP-tagged Smb1 with myc-tagged Mtr10 is shown. Aco1 served as a negative control. *n=3*

It is currently unclear whether the Yku-heterodimer is loaded in the cytoplasm. The Yku70-Yku80 heterodimer binds to a 48 nucleotide long stem loop of *TLC1* and is required for the nuclear localization of this RNA ^18, 28, 45^ and (Fig. 2A). Therefore, we investigated a potential binding between Mtr10 and Yku70 in co-IP experiments, but we did not detect an interaction (Fig. 4C).

Finally, we investigated, whether Mtr10 might, like Cse1, also interacts with the Sm-ring in co-IPs and indeed, a specific band of Mtr10 was detectable in the eluates of the Smb1-IPs (Fig. 4D). These findings indicate that both import receptors interact with the Sm-ring to support nuclear import of the matured *TLC1* RNP.

### Processing of *TLC1* occurs in the nucleus after re-import of the protein bound RNP

The longer pre-*TLC1* is trimmed by the exosome up to the Sm-ring, resulting in a ~1150 nucleotide long form ^6^. Our studies in the re-import mutants suggest that trimming occurs after shuttling, because of an increased Smb1-binding to the immature precursor RNA of *TLC1* (Fig. 1I, 2H). Moreover, mutants that accumulate *TLC1* in the cytoplasm accumulate immature *TLC1* at decreasing total *TLC1* levels (Fig. 2I-K), suggesting that the *TLC1* RNA is not trimmed before its journey through the cytoplasm.

To further analyze this point, cytoplasmic fractionation experminets were carried out. Compared to wild type cells, more unprocessed than total *TLC1* accumulates in the cytoplasm of the import factor mutants (Fig, 6A and 6B). These findings suggest that the immature form shuttles into the cytoplasm and trimming occurs afterwards by the nuclear exosome. Secondly, we assayed the *TLC1-*forms in the *mex67-5* nuclear export mutant. We detected a decrease of total *TLC1*, which is in agreement with earlier findings in which Mex67 was suggested to protect *TLC1* from its degradation ^29^. However, in our experiments we still see a two-fold increase of the immature *TLC1* in *mex67-5* mutants, which suggests that immature *TLC1* is not immediately degraded when Mex67 is missing (Fig. 6E). Possibly, other proteins such as the guard proteins, one of which is Npl3, that recruit Mex67 protect the immature form from being degraded while waiting for export. To test whether Npl3 binds *TLC1*, we precipitated the protein and found *TLC1*, in particular the immature form to bind Npl3 (Fig. 6 C, D). To address whether the trimming to the mature *TLC1* occurs via the nuclear exosome, we analyzed *TLC1* in *rrp6Δ*. Its deletion leads to an approximately two-fold increase of the total *TLC1*, which includes the mature form and the immature form, and a ~7-fold increase of the immature form of *TLC1* (Fig. 6E), indicating that the immature form is trimmed in the nucleus. Remarkably, although the RNA is less stable in *mex67-5* mutants, the immature *TLC1* still increased five fold in the double mutant *rrp6*Δ *mex67-5*, suggesting a) that its trimming occurs in the nucleus and b) that the immature form is still protected without Mex67. Furthermore it has to be noted, that in contrast to Vasianovich and colleagues, we find no rescue of the decreased *TLC1* in the double mutant *rrp6*Δ *mex67-5* as detected by northern blot ^29^. Since we applied in our experiments the highly sensitive qPCR method, we were able to detect an increase of the immature *TLC1* when its export is blocked in *mex67-5*, despite the overall decrease of the total *TLC1* in both mutants (Fig. 6E).

**Figure 6.**
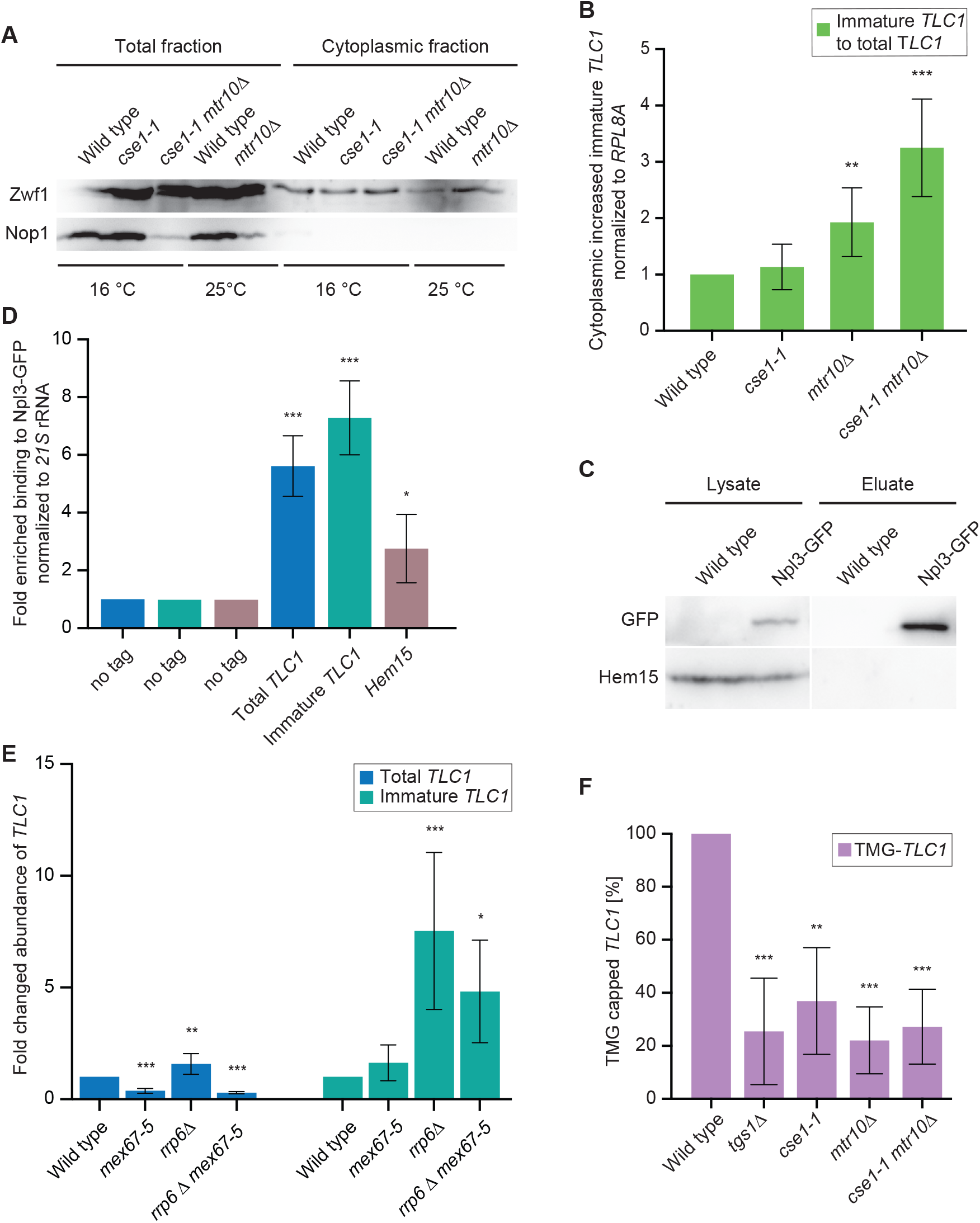
Processing and TMG-capping of *TLC1* occurs in the nucleus upon *TLC1* re-import. (A) Unprocessed *TLC1* accumulates in import factor mutants. A Western blot of a nucleo-cytoplasmic fractionation experiment is shown for the indicated strains. The nuclear Nop1 protein and the cytoplsasmic Zwf1 protein served as controls for successful isolation of the cytoplasmic fraction. Strains were shifted to 16 °C for 1h 15min prior to the experiments. *n =5* (B) qPCRs of the fractionation experiments shown in A reveals an accumulation of the unprocessed *TLC1* in the import factor mutants at the permissive termperature. *n =5* (C) Western blot showing an example of a Npl3 IP used in Fig. 1E in the RIP experiments. (D) *TLC1* binds to Npl3. The Npl3 co-precipitated RNA was used as a template in qPCRs for *TLC1* and Hem15 as positive control. *n = 4* (E) Trimming of *TLC1* is mediated by the nuclear exosome after nuclear re-import. The RNA was isolated from the indicated strains after shifting them to 37°C for 2h. Subsequently, qPCRs were carried out that determined the amount of the total *TLC1* and its immature, longer precursor. *Mex67-5 and rrp6Δ mex67-5 n =4. Rrp6Δ n =8* (F) TMG-capping occurs after nuclear *TLC1* re-import. The amount of TMG-capped *TLC1* was determined by TMG-co-IPs and subsequent qPCRs from the indicated strains. *tgs1Δ* served as a positive control and the black line indicates the amount of the precipitated non-TMG-capped RNAs resembling the base line for TMG-capping. *Tgs1Δ* and *mtr10Δ n =4. Cse1-1* and *cse1-1 mtr10Δ n = 3.*

Taken together, these finding show that trimming of the immature *TLC1* requires nuclear export and that correct trimming occurs only after Sm-ring loading, which seems logic, because the Sm-ring was shown to protect *TLC1* from full degradation ^16^. Our data furthermore suggest that the immature *TLC1* is protected from degradation by Mex67 binding and its immediate export activity into the cytoplasm. But they also suggest that additional proteins, like Npl3, protect the immature *TLC1* from degradation, possibly the guard proteins that recruit Mex67 after quality control, as the immature form increases in the nucleus of *mex67-5* mutants (Fig. 6D and 6E). Our data further indicate that rather the mature form of *TLC1* is degraded when Mex67 is mutated, which might reflect its regulsar turnover rate when no new telomerase is made.

Another modification of *TLC1* is the TMG-capping at its 5’ end in the nucleolus by Tgs1 ^10, 46^. To investigate, whether this trimethylation occurs before or after shuttling we carried out RIPs experiments with a TMG-cap specific antibody. Subsequently, we carried out qPCRs to detect *TLC1*. As compared to wild type, we found an approximately 30% decrease in the level of the TMG-capped *TLC1* in a *TGS1* deletion strain, which resembles the zero-line for the unspecific binding of this antibody, because Tgs1 is the only trimethyltransferase in yeast. Clearly, also in the single import mutants *mtr10*Δ and *cse1-1* as well as in the double mutant we found a reduction, which was around 30% (Fig. 6F). These findings indicate that TMG-capping occurs after shuttling.

Taken together, we have uncovered a stepwise maturation cycle for the telomerase, which is summarized in Fig. 7. Maturation begins with the nuclear export of the longer precursor *TLC1* into the cytoplasm by Mex67, probably assisted by Npl3 and Xpo1 that binds to the CBC. In the cytoplasm not only the Sm-ring is loaded, but also the Est- and the Pop-proteins attach to the immature *TLC1*. Its successful assembly allows Mtr10 and Cse1 to bind and re-import pre-*TLC1* into the nucleus. Subsequently, trimming of the immature form into the mature ~1150 nucleotide long form is mediated by the nuclear exosome and TMG capping finally occurs in the nucleolus by Tgs1.

**Figure 7.**
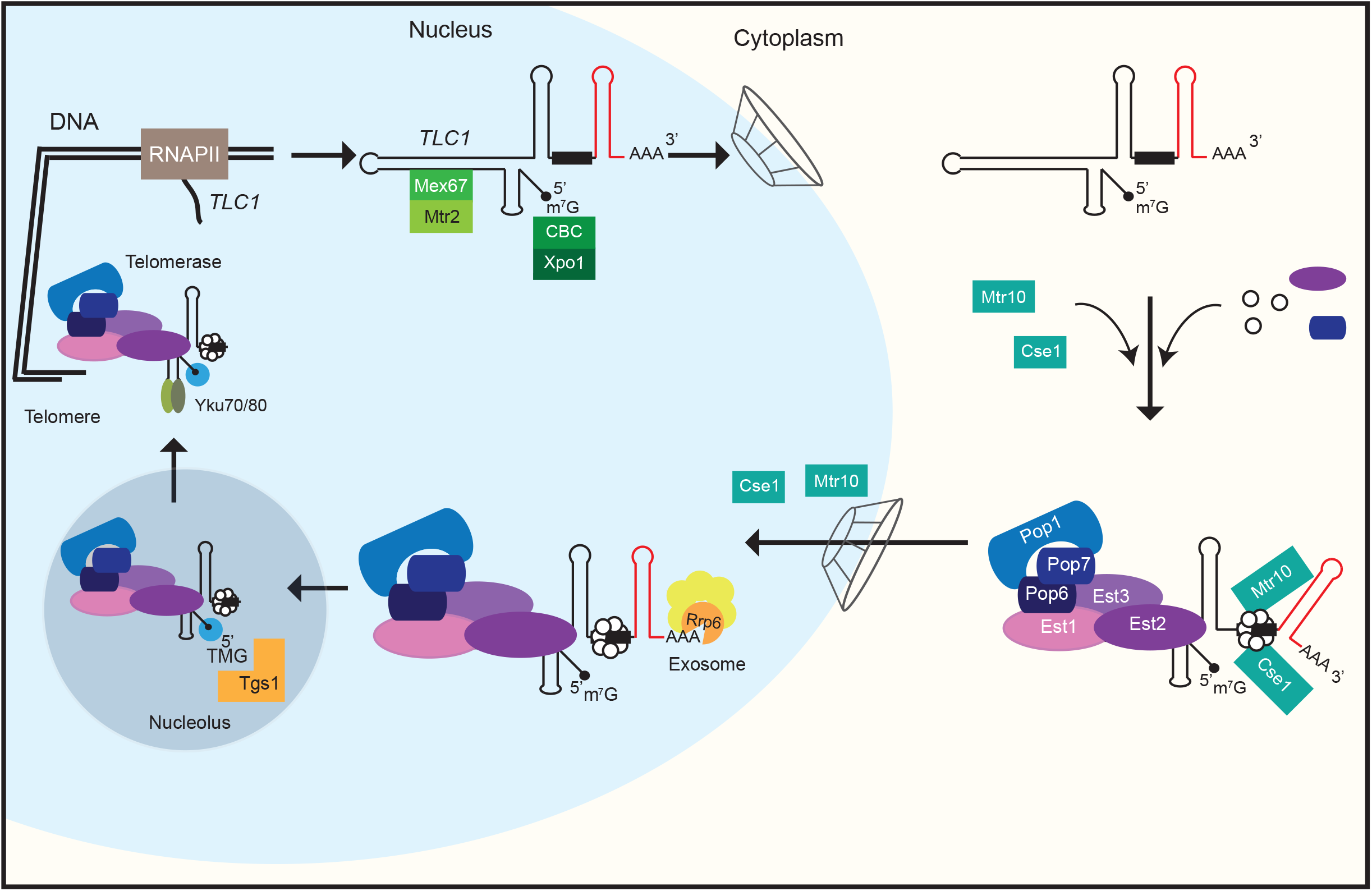
Model for the compartmental stepwise maturation of *TLC1*. The longer precursor of *TLC1* is transcribed in the nucleus and immediately exported into the cytoplasm upon binding of the export receptor Mex67-Mtr2 and the karyopherin Xpo1/Crm1, the latter of which interacts with CBC bound m^7^G-caps. Upon export, Mex67 and Xpo1/Crm1 are displaced and the Sm-ring, the Est- and the Pop-proteins assemble on the immature *TLC1* in the cytoplasm. Subsequently, this RNP is re-imported back into the nucleus via Mtr10 and Cse1, both of which contact *TLC1* via its Sm-ring. Thus, nuclear re-import can only occur after RNP formation including the Sm-ring and therefore resembles an important quality control checkpoint. In the nucleus, the import receptors dissociate and *TLC1* is trimmed up to the Sm-ring by the nuclear exosome. Finally TMG-capping occurs in the nucleolus assisted by Smb1, which interacts with Tgs1. This step terminates shuttling, because export receptors cannot be loaded anymore. The matured holoenzyme subsequently acts in telomere maintenance.

In our compiled investigation we have ordered the maturation process of *TLC1* and show the order of events from nuclear export of the premature transcript to the binding of the Est- and Pop-proteins and the Sm-ring in the cytoplasm to its trimming and TMG-capping after re-import into the nucleus. We show that the Sm-ring binding resembles a quality control step that enables subsequent movement of the RNP into the nucleus. Thus, the step-wise maturation of *TLC1* is controlled by its compartmental shuttling.

## Discussion

Functional telomerases are important to prevent the repetitive shortening of the DNA ends after each round of replication for recurrent cell divisions. Due to the fact that these holoenzymes undergo a stepwise assembly, a danger is that incompletely assembled telomerases interfere with the process of telomere elongation. In the worst case, these immature telomerases could, due to their restricted or absent functionality, prevent telomere maintenance. To avoid such a scenario, a mechanism has developed, in which the RNA scaffold, *TLC1* in yeast, is exported into the cytoplasm via the export receptors Mex67-Mtr2 and Xpo1/Crm1 ^23, 28^. While Xpo1/Crm1 contacts the CBC, which is bound to the monomethyl cap of the RNA-polymerase II transcribed RNA, Mex67-Mtr2 was speculated to contact the RNA via the adapter proteins Npl3, Gbp2 and Hrb1, because such contact is observed for mRNA and snRNA export ^24, 25, 26, 27^.

Binding of the Est proteins to *TLC1* and association of the Sm-ring were suggested to be cytoplasmic events, while TMG-capping and trimming of *TLC1* were suggested to occur in the nucleus prior to shuttling ^23, 28, 29^. Up to date, the place of the Pop-protein loading onto *TLC1* was unknown, but suspected to occur in the cytoplasm, because mutants of *POP1* and *POP6* accumulated *TLC1* in the cytoplasm ^20^. We collected additional evidence for a cytoplasmic Pop-protein loading and analyzed the localization of Pop1 in the import factor mutants. We found an accumulation of Pop1 in the mutants (Fig. 3C), supporting the idea of their cytoplasmic loading and supporting the newly discovered role of Cse1 in *TLC1* nuclear import. Finally, we could confirm this result through RIP experiments in which we detected a decreased binding to Pop1 in *mex67-5* and an increased binding in the import mutant *cse1-1* (Fig. 3E, G). As the function of the Pop-proteins is to stabilize the Est proteins on the RNA and to support the RNP structure ^21, 22^ their cytoplasmic loading seems to be a final step before re-import.

Another important event is the loading of the Sm-ring that limits the trimming of the *TLC1* precursor, which was recently shown to occur in the cytoplasm, similar to the Sm-ring loading onto snRNAs ^24, 29^. However, in the interesting study of Vasianovich and colleagues a modified *TLC1* that contained several MS2 stem loops was used, which could have an influence on its localization. In our study we used probes and primers that 1) detect the unmodified natural *TLC1* and 2) selectively detect the immature *TLC1*-precursor. With this method we confirmed the data of Vasianoivich and colleagues and show that the Sm-ring loading is a cytoplasmic event. But in addition, we have shown that the Sm-ring and the Pop-proteins associate with the immature precursor of *TLC1* in the cytoplasm (Fig. 1I, 2H and 3G) and that trimming occurs after Sm-ring loading and after re-import into the nucleus (Fig. 1I, 1K, 2H-K, 3E, 3G, 6E).

It was shown that the karyopherin Cse1 contacts the Sm-ring and particularly Smb1 for nuclear re-import of the snRNAs ^24^. Interestingly, we also detected a binding between the Sm-ring and the other import factor, Mtr10 (Fig. 4D). This finding is important for three reasons. First, besides Mtr10, Cse1 is another import factor that facilitates the transit of this huge RNP through the NPC. The more transport factors are involved in transporting large particles, the better the transit is working. This is supported by the additive effect for the mislocalization of *TLC1* seen in the combined import factor mutants (Fig. 6A,B). Secondly, a potential failure of one of the import receptors does not fully prevent *TLC1* re-import. Third, nuclear re-import supported by Mtr10 and Cse1 can only occur on *TLC1* molecules that have received the Sm-ring, which resembles an elegant way of cellular quality control for this important maturation step of the telomerase. For this, Cse1 might be more important because we detect a decreased binding of *TLC1* to Smb1 in *cse1-1* cells, even though we see no drastic change in the overall amount of *TLC1* (Fig. 2F-K). Together, these findings argue for a control step in the RNP formation, which prevents that faulty RNPs can enter the nucleus and jeopardize telomere function. However, we cannot exclude that another protein is involved in its stabilization, which is dependent on Cse1 transport. In every sense, Sm-ring loading is an essential step in maturation and allows nuclear re-import of pre-*TLC1*.

Interfering with the shuttling of *TLC1* either by mutation of the export factors Mex67 and Xpo1 or the import receptor Mtr10 was shown to result in telomere shortening defects ^23, 30^. Interestingly, we did not detect any telomere shortening defects for the novel *TLC1* re-import factor Cse1. Instead, mutation of *CSE1* resulted in a Type I like survivor phenotype, which is characterized of being capable to amplify subtelomeric Y’ elements (Fig. 5A, B). Type I survivors can escape cell death by lengthening their telomeres through homologous recombination ^37^. *cse1-1* seems to belong to the Type I survivors, because it shows the typical highly amplified subtelomeric Y’ elements generated via homologous recombination, even though *cse1-1* cells show no shortening in the termial fragment. Thus, while the export receptor mutants and *mtr10*Δ did not allow telomere elongation via recombination, a mutation in *CSE1* does. As the Y-element amplification is based on homologous recombination by the recombinase Rad51, one can speculate that either Rad51 itself, or factors of the Rad51 homologous recombination pathway might not be imported in *mtr10*Δ, so that *mtr10*Δ mutants cannot use homologous recombination for telomere maintanace but rather show a shortening of the telomeres. Other survivors are mutants of telomere capping components, since very short or uncapped telomeres are prone to recombination despite a functional telomerase ^47, 48, 49^. Therefore, we cannot exclude that Cse1 might import one of these proteins. Nevertheless, the mislocalization of *TLC1* in the cytoplasm of the *cse1* mutants suggests rather a direct effect.

Moreover we have excluded an involvement of Srp1 in the localization of *TLC1*. In the *srp1-31* mutant, no mislocalization of *TLC1* was observed (Fig 5D). Srp1 was suggested to mediate the nuclear localization of Est1 and since Srp1 mislocalizes in the *cse1-1* mutant an influence was conceivable ^41, 42^. However, the type I like survivor phenotype does not appear to be dependent on Srp1, as mutation of *SRP1* does not lead to a mislocalization of *TLC1*.

At steady state about 80 to 90 % of *TLC1* are present in the mature shorter form which is present in the functional telomerase and only a small fraction is present as the polyadenylated longer form ^12, 8^. Older model discuss the CPF-CF terminated poly(A)^+^ form as a precursor of the mature form ^12^, but more recent studies suggest that the NNS terminated form might be the prominent one ^14^. Therefore, it is currently unclear how much of the *TLC1* originates from NNS or CPF-CF termination. However, since both versions a) need to adopt the Sm-ring for general protection, which was shown to be a cytoplasmic process ^24^ and b) total *TLC1* significantly decreases if shuttling is prevented (Fig. 2F, I) and c) leads to a defect in the telomeres (Fig. 5) and ^23^, shuttling of any immature form seems highly relevant.

It was unclear when trimming occurs. Earlier studies indicated that mutations in the nuclear component of the exosome resulted in a strong increase of *TLC1* ^16^. However, the experimental set up neither allowed to distinguish between the mature form or the immature poly(A)^+^ precursor of *TLC1*, nor whether the exosomal trimming occurred before or after shuttling. To determine the time and place of the *TLC1* trimming, we first showed that the immature form accumulates in nuclear re-import mutants and in mutants that are defective in the Sm-ring assembly (Fig. 1 and 2I-K). Additionally we have shown by nucleo-cytoplasmic fractionation experiments that the immature form accumulates in cytoplasm in the import factor mutants (Fig 6B), which indicates that trimming occurs after re-import. Furthermore, mutation of *RRP6* resulted in a strong increase of the immature *TLC1* precursor (Fig. 6E), suggesting that the nuclear exosome is responsible for trimming after shuttling. This finding also suggests that the immature form is still protected without Mex67. One possibility is that the Mex67-interacting and RNA-binding proteins Npl3, Gbp2 and Hrp1 might guard *TLC1*. And indeed, we have shown that *TLC1* and especially the immature *TLC1* is bound by Npl3 (Fig. 6D). Co-transcriptionally loading of Npl3 prior to binding of Mex67 ^50, 51^ provides coverage and thus protection of the RNP.

The finding, that Sm-ring binding occurs before trimming seems logical, because unprotected RNA is an ideal substrate for the exosome ^17^. Interestingly, preventing nuclear export by mutation of *MEX67* leads to a decrease of *TLC1* (Fig. 6E and ^29^). During the short period of time in which the immature *TLC1* RNA is usually present in the nucleus before Mex67 and Xpo1 export it into the cytoplasm, the nuclear exosome is not able to attack the RNA sustainably. Thus, Mex67 cannot be the only protecting RNA-binding protein, because even in mutants of this export receptor we detect a ~5-fold increase of the immature *TLC1* in *rrp6*Δ *mex67-5* as compared to wild type. Possibly this is achieved by guard proteins such as Npl3 that recruit Mex67 ^27, 51^.

Our finding that the Sm-ring loading occurs in the cytoplasm prior to TMG-capping is in agreement with other studies that have shown that the Sm-ring is crucial for trimethylation of the cap ^52^. Furthermore, we have shown earlier that snRNAs are retained in the nucleolus when Smb1 is depleted that showed that an intact Sm-ring is required for efficient trimethylation and subsequent nucleolar release into the nucleoplasm. It is well conceivable that the formed Sm-ring assists TMG-capping, as it interacts with Tgs1 both, *in vivo* and *in vitro* ^10, 24^.

Importantly, snRNA nucleo-cytoplasmic shuttling was suggested to be terminated by trimethylation of the 5’ cap ^24^. The export receptor Xpo1 interacts with the 5’ cap binding complex CBC, in particular with Cbp80 ^24^. This might be similar for *TLC1*, because pre-*TLC1* contains an m7G cap, which is bound to CBC (Fig. 1K) and *TLC1* interacts with Xpo1 ^23^. After receiving the TMG cap, CBC might not be bound and Xpo1 then unable to interact. Since the other export receptor, Mex67, is already displaced at the NPC when entering the cytoplasm ^53, 54^, Xpo1 is the only transport factor left for a repeated snRNA export and TMG capping could close this option. Therefore, it seems logic that TMG capping on *TLC1* also occurs after shuttling and maturation and we could indeed show that preventing nuclear re-import of *TLC1* results in the accumulation of m^7^G, but not TMG capped RNAs (Fig. 6F).

Together, our analysis uncovered the steps in which telomerase maturation takes place (Fig. 7) and identified that proper cytoplasmic assembly of the telomerase is prerequisite for nuclear re-import and thus represents a quality control check point in the life cycle of this holoenzyme. Any way of interfering with the compartmental maturation process inevitably leads to defects in telomere biology (Fig. 5) and ^18, 20, 23, 28, 30, 55, 56^, reflecting the importance of this step-wise process.

## Experimental procedures

### Yeast strains, plasmids and oligonucleotides

All yeast strains used in this study are listed in the Supplemental Table 1, oligonucleotides in Supplemental Table 2 and plasmids in Supplemental Table 3. Plasmids and yeast strains were generated by conventional methods.

### Fluorescent *in situ* hybridization experiments (FISH)

The experiments were essentially carried out as described ^24^. RNA probes were with Cy3-labeled oligonucleotides (Sigma), which are listed in Supplementary Table 2. Cells were grown to mid log phase (1×10^7^ cells/ml) prior to temperature shift to 37 °C 1h or 2h or to 16 °C for 1 h 15 min (Fig. 2A, 5D). For Sm-ring dependent localization studies, cells were grown to log phase in YP medium containing 2 % galactose. Afterwards 4 % glucose was added and cells were incubated at 25 °C for 2 h (Fig. 1E). Samples were fixed by adding formaldehyde to a final concentration of 4 % for 45 min at room temperature. Cells were spheroplasted by adding zymoylase, subsequently permeabilized in 0.1 M potassium phosphate buffer pH 6.5, 1.2 M sorbitol, 0.5 % Triton® X-100, pre-hybridized with Hybmix (50 % deionized formamide, 5× SSC, 1x Denhardts, 500 μg/ml tRNA, 500 μg/ml salmon sperm DNA, 50 μg/ml heparin, 2.5 mM EDTA pH 8.0, 0.1 % Tween® 20) for 1 h on a polylysine coated slide at 37 °C and hybridized in Hybmix with the specific probe over night at 37 °C. After hybridization, cells were washed with 2x SSC and 1x SSC at room temperature, each for 1 h and 0.5x SSC at 37 °C and room temperature, each for 30 min. DNA was stained with Hoechst 33342 (Sigma). Microscopy studies were performed with a Leica AF6000 microscope and pictures were obtained by using the LEICA DFC360FX camera and the LAS AF 2.7.3.9 software (Leica). For deconvolution (Fig. 1E, 2A and B) z-stacks (10 stacks; 0,2 *μ*m) were recorded and the maximal projection was deconvoluted with 3 iterations by the LAS AF 2.7.3.9 software (Leica).

### RNA co-immunoprecipitation experiments (RIP)

All yeast strains were grown to mid log phase (2×10^7^ cells/ml). For RIP experiments seen in Fig. 1 A, B, J, K; 2 C, D and 6 C and D cells were cultured at 25 °C, in 1 C, D, H, I; 2E-H; 3D-G and 6F cells were shifted to a non-permissive temperature for 1 h 37 °C or 16 °C respectively. Afterwards cells were harvested and lysed in RIP buffer (25 mM Tris HCl pH 7.5, 100 mM KCl, 0.2 % (v/v) Triton X-100 (1% Triton X-100 for 2E), 0.2 mM PMSF, 5 mM DTT, 10 U RiboLock RNase Inhibitor (Thermo Scientific) and protease inhibitor (Roche) using the FastPrep^®^-24 machine (MP Biomedicals) three times for 30 sec at 5.5 m/s. After centrifugation the supernatant was incubated for 1 h at 4 °C with GFP-Selector beads (NanoTag) (Fig. 1 B,D, H, I, K; 2 D, F-H; 3 E, G and 6D). For TMG-cap-IPs total RNA was extracted from yeast lysates using trizol-chloroform (Ambion^®^ RNA by Life technologies™). 50 *μ*g of the total RNA was incubated for 1 h at 4 °C with 10 *μ*l of the anti-2,2,7-trimethylguanosine-antibody (Calbiochem Milipore) coupled to sepharose beads (Fig. 6F). Detection occurred via an anti-digoxygenin antibody coupled to alkaline phosphatase (1:10,000) (Roche).

The beads were washed five times with RIP buffer and for GFP-RIP split in two portions after the last washing step. Proteins were detected by western blot (1A, C, G and J; 2C, E; 3 D, F and 6C). Eluates were purified via trizol-chloroform (Ambion^®^ RNA by Life technologies™) extraction. The purified RNA was measured via Nanodrop and a defined amount of RNA was reverse transcribed with FastGene Scriptase II (Nippon Genetics) for subsequent qPCR analyses. To normalize the total RNA amount, we used the mitochondrial 21S rRNA. For comparisons, we always related the immature to the total *TLC1*. All eluates were related in each strain to the lysates to account for strain-specific variations in the total amount.

### Total RNA Isolation

Total RNA Isolation was carried out with the NucleoSpin RNA Kit from Macherey-Nagel. All steps were performed according to the manufactures description, except step 7. The DNA digestion on the column was executed for 1 hour. An additional DNA digestion step was carried out after elution of the RNA. For the additional DNA digest the eluted RNA was mixed with a 10th volume of the Reaction Buffer for rDNAse and 1 *μ*l rDNAse, according to the manufactures description. The digest was incubated for 1 hour at 37°C, before it was terminated through sodium acetate ethanol precipitation. For this 0.1 volume 3 M sodium acetate, pH 5.2, 2.5 volumes of 99 % pure ethanol and 1 *μ*l glycoblue were mixed with the RNA and incubated over night at – 20 °C. For normalization, the 21S rRNA amount was determined. In Fig. 2H and K and 6B, the enrichment of the unprocessed form was shown in comparison to total *TLC1* amount. For this purpose, the total amount present in each strain was used as normalization to account for strain internal variations.

### Nucleo-cytoplasmic fractionation experiment

For the detection of *TLC1* in the cytoplasm (Fig. 6B) cells were grown to mid log-phase (2×10^7^ cells/ml). The cells were harvestet by centrifugation for 5 min at 4000 rpm. Cells were washed once with 1ml YPD/ 1 M Sorbitol/ 2 mM DTT and resuspended in YPD/ 1 M Sorbitol/ 1 mM DTT. Cells were spheroblasted using 1 mg zymolyase (100 mg/ml) and after that diluted in 50 mL YPD/ 1 M Sorbitol for 30 min at 25°C for recovery. Subsequently, the cells were shifted to 16°C for 1 h 15min (*cse1-1* and *cse1-1 mtr10Δ)*. Cells were placed on ice, centrifuged at 2000 rpm for 10 min and the pelleted cells were resuspended in 500 μl Ficoll buffer (18% Ficoll 400, 10 mM HEPES pH 6.0) and 1*μ*l Ribolock. Cells were lysed by addition of 1 ml buffer A (50 mM NaCl, 1 mM MgCl2, 10 mM HEPES pH 6.0). The suspension was mixed and centrifuged at 4000 rpm for 15 min. The supernatant was used for cytoplasmic analyses. RNA was isolated using the Nucleo-Spin RNA Kit (Macherey and Nagel). The purified RNA was reverse transcribed with FastGene Scriptase II (Nippon Genetics) for subsequent qPCR analyses. All values were normalized to the amount of the mRNA present for *RPL8A*. To verify no nuclear contamination in the cytoplasmic fraction, aliquots of the samples were analyzed in western blots for the presence of the cytoplasmic Zwf1 protein and the absence of the nuclear Nop1 protein (Fig. 6A).

### GFP-microscopy

Cells were grown, treated and harvested as described in the FISH experiments. Cells were fixed with 3 % formaldehyde for 2 min at room temperature, subsequently washed with 0.1 M phosphate buffer pH 6.5 and with P-solution (0.1 M phosphate buffer pH 6.5, 1.2M Sorbitol), before an aliquot was added to a polylysine-coated slide for 30 min at 4 °C. Permeabilization of the cells, DNA staining and microscopy was performed as described in the FISH experiment.

### Co-immunoprecipitation (IP) experiments

All yeast strains were grown to log phase (2-3×10^7^ cells/ml). Afterwards, the cells were harvested and lysed in IP buffer (1 x PBS, 3 mM KCl, 2.5 mM MgCl_2_, 0,5 % Triton X-100 and protease inhibitors from Roche). 35 *μ*l of this lysate was loaded onto an SDS-gel (lysate lanes). The supernatant was incubated for 1 h at 4 °C with GFP-Selector beads (NanoTag) (Fig. 4A, B and D) or with Myc-trap beads (Chromotek) (Fig. 4C). The beads were washed five times with IP buffer, and finally resuspended in 35*μ*l SDS-sample buffer. The entire eluate sample was loaded onto the SDS-gels.

Subsequently, the proteins were detected by Western blot analyses with the indicated antibodies (GFP (Chromotek) 1:4,000; c-myc (9E10) (Santa Cruz) 1:1,000; Hem15 and Grx4 each 1:5,000 and Aco1 1:2,000 (U. Mühlenhoff); Nop1 (Santa Cruz) 1:4,000; Hdf1(Yku70) (Santa Cruz) 1:4,000). Signals were detected with the Fusion SL system (PeqLab) and FusionFX7 Edge (Fusion FX Vilber). To be able to detect several proteins in one experiment, the western blots were cut horizontally according to the size of the desired proteins to be able to detect each stripe with individual antibodies

### Drop dilution analysis

Cells were grown to log phase (2-3×10^7^ cells/ml) and diluted to 1×10^7^ cells/ml. 10-fold serial dilutions to 1×10^3^ cells/ml were prepared and 10 *μ*l of each dilution was spotted onto full medium (YPD) agar plates that were subsequently incubated for 3 days at the indicated temperatures (Fig. 2 B). Pictures were taken after 2 and 3 days with the Intelli Scan 1600 (Quanto technology) and the SilverFast Ai program.

### Southern blot analysis

Yeast strains were either passaged in liquid culture (Fig. 5B) or on solid agar plates (Fig. 5C). For passaging in liquid culture, cells were inoculated with a starting concentration of 1×10^5^ cells/ml. The strains were grown for 3 days per passage at 20 °C until they reached 1×10^8^ cells/ml and were afterwards diluted again to 1×10^5^. Overall 8 passages, corresponding to 80 generations, were carried out.

Cell passages on solid agar plates were used for generating Type I survivors in *tlc1Δ* cells. The strains were freshly restreaked form the database and single colonies were restreaked after 2-3 days at 25°C. One restreak comprises about 25 generations. Overall 5 restreaks were generated coresponding to about 125 generations.

Genomic DNA was isolated with the MasterPure Yeast DNA Purification Kit (Lucigen). This gDNA was digested with XhoI (Nippon Genetics) according to the manufactures description. 20 *μ*g gDNA were loaded per lane and electrophoretically separated on a 1 % TBE gel at 4°C for 3 hours. The gel was depurinated (250 mM HCl) for 15 minutes, denatured (1.5 M NaCl, 0.5 M NaOH) for 30 minutes, neutralized (1.5 M NaCl, 0.5 M Tris-HCL, pH 7.5) for 30 minutes and equilibrated in 20x SSC (0.3 M Tri-sodium citrate, 3 M NaCl, pH 7.0) for 15 minutes. A capillary blot was used for the transfer onto a HybondN^+^ (GE Healthcare) membrane. After transfer, the membrane was crosslinked for 7 min by UV light (254 nm, 120000 *μ*J/cm^2^) and then heated to 80 °C for 2 h. Subsequently, pre-hybridization was carried out for 1h at 68 °C in pre-hybridization buffer (0.5 M sodium phosphate buffer pH 7,5; 7% (w/v) SDS; 1 mM EDTA). For hybridization, a digoxygenin-labeled probe, detecting the TG repeats of the telomeres, was added to the pre-hybridization solution and incubated over night at 37°C with rotation. After washing, the southern blot was detected by using an anti-digoxygenin antibody coupled to alkaline phosphatase (Roche) and the labeling and detection starter kit (Roche).

### Quantification

All experiments shown in this work were performed in biologically independent repetitions as indicated in the figure legends. Error bars represent the standard deviation. P values were calculated using a two-tailed, two-sample equal variance t-test. P values are indicated as follows: ***p < 0.001, **p < 0.01, *p < 0.05.

## Acknowledgements

We are grateful to R. Bordonné, E. Hurt, U. Mühlenhoff, R. Lill, V. Lundblad, E. O’Shea, P.A. Silver, A. Tartakoff and K. Weis for providing plasmids, strains or antibodies. This work was supported from the Deutsche Forschungsgemeinschaft (DFG) and the SFB860 awarded to H.K.

## Author contributions

Experiments were designed and data interpreted by A.G.H., D.B., J-P.L. and H.K.; all experiments were performed by A.G.H. except 1 E, which was done by D.B; 2A by A.G.H. and D.B.; 2B, 3A,B,C by A.G.H and J.-P.L.. The manuscript was written by H.K.; all authors discussed the results and commented on the manuscript.

## Supplementary information to

### Supplementary Tables

**Supplemental Table 1.**
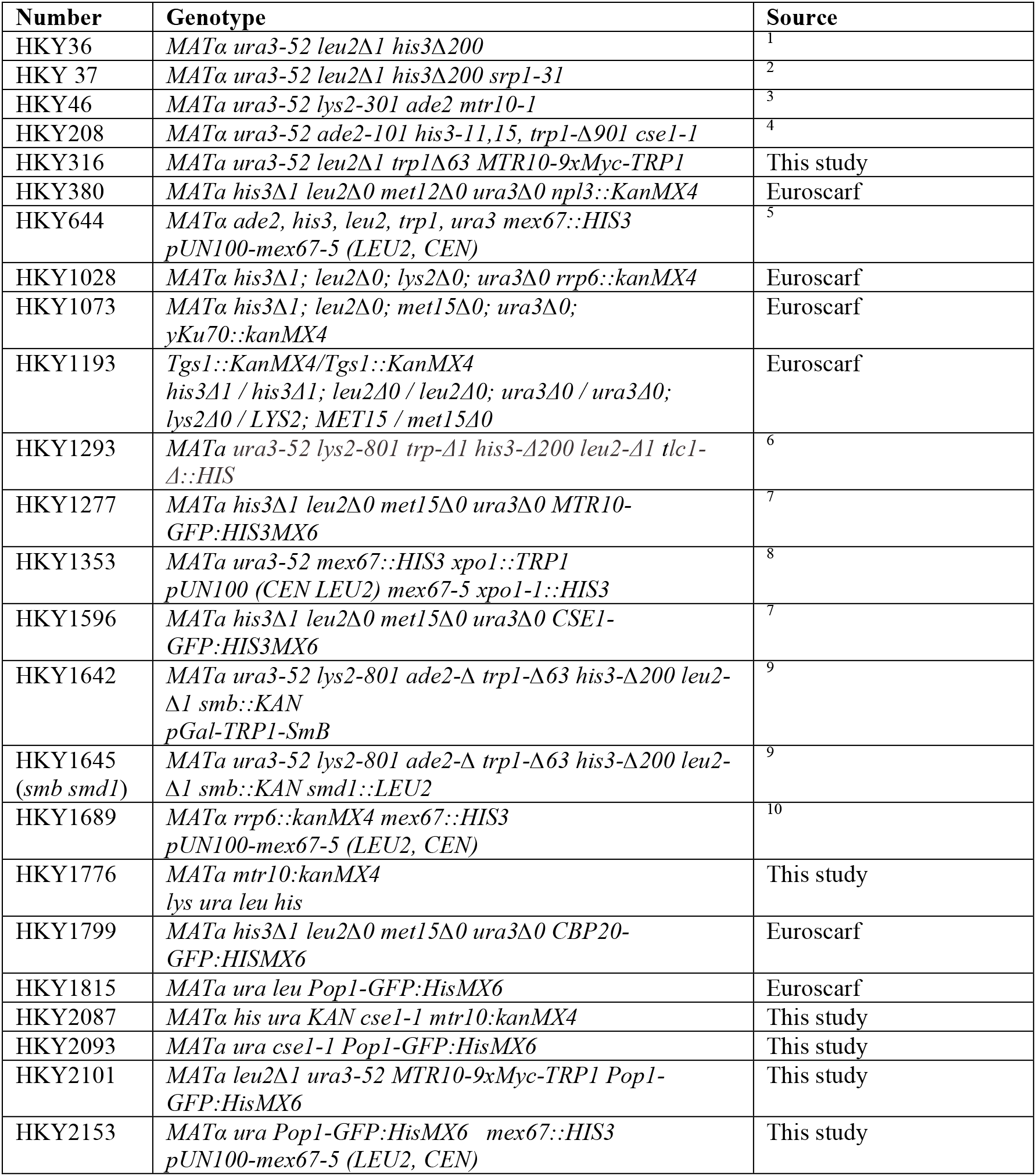
Yeast strains used in this study. Related to Figures 1–6

**Supplemental Table 2.** Plasmids used in this study. Related to Figures 1–6.

| Number | Genotype | Name |
| --- | --- | --- |
| pHK88 | CEN URA3 | 11 |
| pHK206 | CEN URA CSE1 | This study |
| pHK765 | CEN URA GFP-Npl3 | 12 |
| pHK1469 | CEN URA SmB-GFP | 13 |
| pHK1483 | CEN URA GFP-POP1 | 14 |
| pHK1589 | URA3 EST1-(Gly)6-(myc)12 | 15 |
| pHK1606 | CEN URA pAdh-Est1-GFP | This study |

**Supplemental Table 3.**
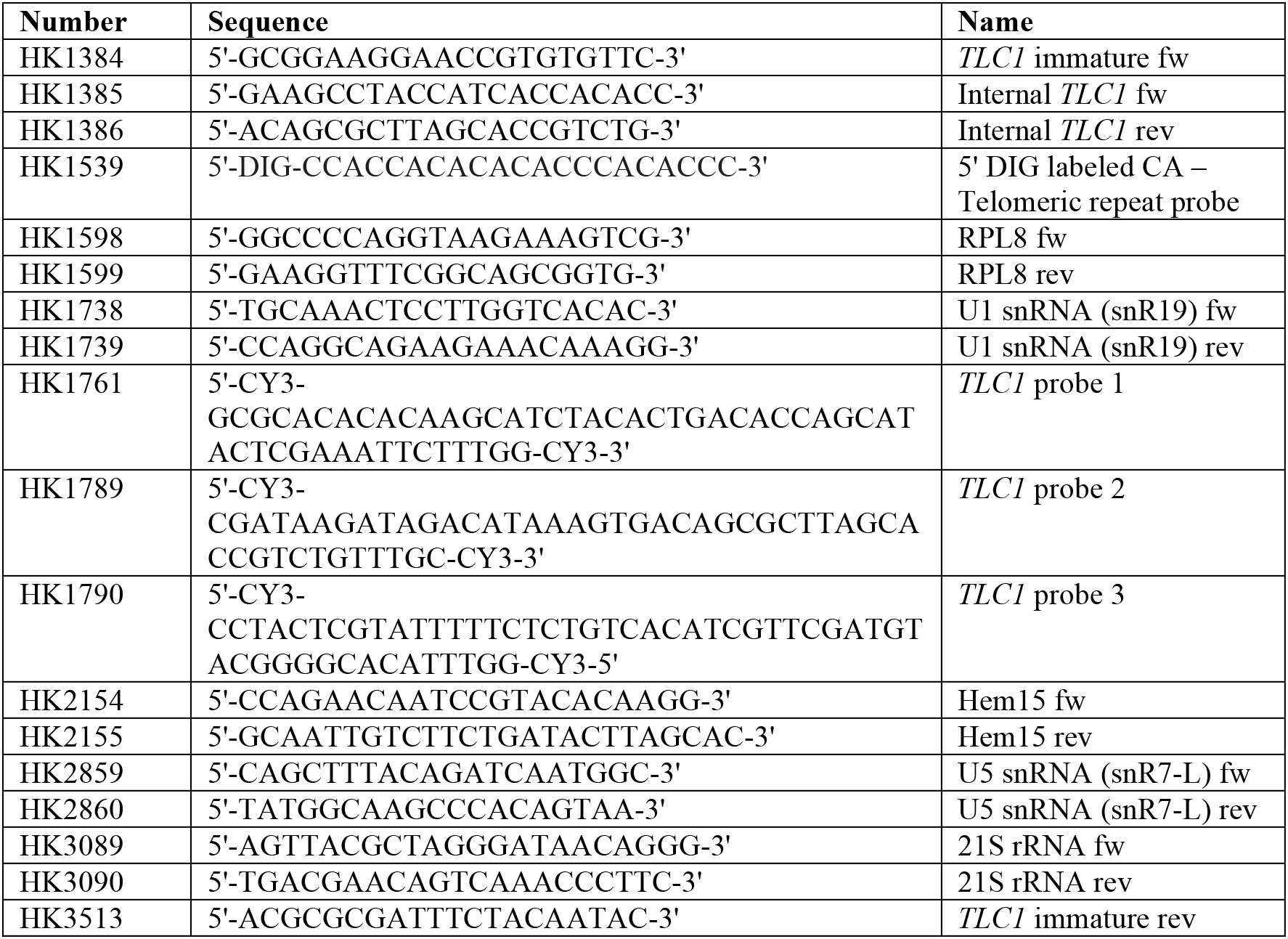
Oligonucleotides used in this study. Related to Figures 1–3, 5, 6. Forward primer (fw) and reverse primer (rev).

